# Contribution of private gardens to habitat availability, connectivity and conservation of the common pipistrelle in Paris

**DOI:** 10.1101/579227

**Authors:** Anne Mimet, Christian Kerbiriou, Laurent Simon, Jean-François Julien, Richard Raymond

## Abstract

Urban sprawl is one of the greatest global changes with major negative impacts on biodiversity and human well-being. Recent policies have acknowledged the value of urban green areas in counterbalancing such impacts. These policies aim to increase the ecological value of green areas, making cities more permeable to natural populations. However, they are largely focused on the role and management of public green areas, ignoring the role and potential of private green areas for urban ecological value.

This study aims to evaluate the benefits of considering private green areas for conservation efforts in cities. Using data on bat activity and information on vegetation and building height, we quantify the respective role of public and private green areas in habitat availability and connectivity for the common pipistrelle in the city of Paris, France. Our results show that despite the low proportion of private green areas in Paris (36% of the total green areas), they still contributed up to 47.9% of bat habitat availability and decrease the resistance of the city matrix by 88%. The distribution in the city matrix and vegetation composition of those areas appeared especially beneficial for bat habitat availability and connectivity. The study demonstrates the importance of private green areas in the ecological value of cities in complementing the role of public green areas. Our results confirm the need to develop more inclusive urban conservation strategies that include both public and private stakeholders.

**Highlights:**

- The urban ecological value of private gardens outweighs that of public gardens
- This is true for both habitat availability and connectivity
- Biodiversity policies in cities should also focus on private green areas
- Inclusive conservation strategies are also needed in cities

## 1. Introduction

Urbanization is one of the main driver of the biodiversity crisis, leading to the erosion of species diversity (Mcdonald, Kareiva, & Forman, 2008; Olden, Poff, & McKinney, 2006), a decrease in both total community abundance (Newbold et al., 2015; V. Pellissier, Mimet, Fontaine, Svenning, & Couvet, 2017) and in the abundance of most species (C. G. Threlfall, Law, & Banks, 2012), and to biotic homogenization (McKinney, 2006). The underlying mechanisms include direct loss and fragmentation of natural habitat (Devictor et al., 2008; Devictor, Julliard, Couvet, Lee, & Jiguet, 2007), as well as disconnection of the habitat patches of natural populations, thus impeding movement (Clauzel, Jeliazkov, & Mimet, 2018; Tannier, Bourgeois, Houot, & Foltête, 2016).

Depending on their size, composition, configuration and management, urban green areas have the potential to support wildlife populations by providing habitat and by contributing to the connectivity of natural populations (Alberti, 2005; Muratet & Fontaine, 2015; Muratet, Machon, Jiguet, Moret, & Porcher, 2007; Vincent Pellissier, Cohen, Boulay, & Clergeau, 2012; Politi Bertoncini, Machon, Pavoine, & Muratet, 2012; Shwartz, Turbé, Julliard, Simon, & Prévot, 2014). There is increasing recognition of the ecological value of urban green areas is increasing (Breuste, Niemelä, & Snep, 2008; Goddard, Dougill, & Benton, 2010), along with a call to develop a better understanding of the roles of urban green areas in biodiversity in order to guide conservation actions within urban areas (Dearborn & Kark, 2010; Shwartz et al., 2014).

The benefit of urban green areas are tracked by urban authorities and policies, which favor different types of actions to promote urban biodiversity such as ecological management (e.g., late mowing), reduction of pesticides and maintaining and developing ecological corridors, e.g., in London, Dublin, Berlin or Paris (City of Berlin, 2012; City of Dublin, 2016; City of London, 2016; Conseil de Paris, 2018; Ville de Paris, 2017). These policies aim to make cities permeable to natural populations, usually by targeting the larger public green areas but overlooking private ones, such as gardens (Evans et al., 2012; Goddard et al., 2010). Yet, gardens provide significant amounts of green areas and resource for wildlife in urban areas (Cameron et al., 2012; Davies et al., 2009), complementing large public green areas such as parks by increasing habitat availability and connectivity (Colding, 2007; Loram et al., 2008; Melles, Glenn, & Martin, 2003; Rudd, Vala, & Schaefer, 2002). However, our understanding of this complementation process remains fragmented. We do not have estimates or a general understanding of the relative contributions of gardens and public green areas to habitat availability (i.e. the amount of habitat available to a species) and connectivity in cities, especially considering that said contributions are expected to be species-dependent (Lepczyk et al., 2017) and configuration-dependent (Goddard et al., 2010). Urban ecology literature has stressed the importance of different scales (i.e. local to the landscape scale) and of vegetation structure and heterogeneity in explaining the distribution of species and diversity in an urban context (Goddard et al., 2010; Goddard, Dougill, & Benton, 2013; Lepczyk et al., 2017; Melles et al., 2003). Getting accurate estimates of the relative contributions of gardens and public areas to habitat availability and connectivity therefore requires multiscale analyses including vegetation structure and heterogeneity.

With the goal of evaluating the benefits of including private green areas (i.e. gardens) in urban conservation initiatives, we quantify the respective contributions of public and private green areas to habitat availability and connectivity for a common urban bat species, the common pipistrelle (P. *pipistrellus),* focusing the analysis on the city of Paris. The latter is engaged in a plan to promote biodiversity and has reinforced its biodiversity conservation policy, limiting pesticide use and promoting biodiversity-friendly management in public green areas (https://www.paris.fr/biodiversite, https://www.paris.fr/actualites/paris-s-engage-pour-le-zero-phyto-6160). By funding this study, the city of Paris is looking for more knowledge about its ecological networks.

The choice of a bat species is justified by bats conservation status in the EU: bats are one of the few strictly protected mammals living within urban environments (they are included in Annex IV Council Directive 92/43/EEC, 1992). In addition, as a long-lived insectivorous species with a slow reproductive rate, bats are considered good indicators of the response of biodiversity to anthropogenic pressure (Jones, Jacobs, Kunz, Wilig, & Racey, 2009). Previous researches showed the detrimental effects of urbanization on bat populations (Azam, Le Viol, Julien, Bas, & Kerbiriou, 2016; Walsh & Harris, 1996) but certain bat species occur in cities, using trees planted along streets and parks as urban substitutes for their natural foraging habitat (Oprea, Mendes, Vieira, & Ditchfield, 2009; C. Threlfall, Law, Penman, & Banks, 2011). Some species use man-made structures such as breeding roosts and can live in cities if other basic requirements like water access are also met (Marnell & Presetnik, 2010; Simon, Huttenbugel, & Smit-Viergutz, 2004). The strength of the impacts of urbanization on bats appears to be context-dependent, i.e., the degree of urbanization, the amount of vegetation remaining and patch connectivity have been shown to largely explain observed distribution patterns (Oprea et al., 2009; C. Threlfall et al., 2011). Among common bat species, *P. pipistrellus* is one of the most abundant in North European urban areas (Gaisler, Zukal, Rehak, & Homolka, 1998; Hale, Fairbrass, Matthews, & Sadler, 2012). It is also the most abundant bat species in Paris (86% of total bat passes recorded in Paris in the French Bat Monitoring Programme, see below), making it a suitable model for studying the contribution of private green areas to habitat availability and connectivity in Paris.

We quantify the respective contributions of public and private green areas to habitat availability and connectivity for the common pipistrelle by comparing its habitat availability and connectivity in a scenario that includes all of the green areas in Paris (the “All green areas” scenario) one that includes only public green areas (the “No private green areas” scenario). In Paris, private green areas are scattered throughout the city between the much larger public green areas following a density gradient going from the city center to the belt (Figure 1). Based on such contrasting spatial patterns, we expect private and public areas to contribute differently to the habitat availability and connectivity of the common pipistrelle. In a first step, we model the abundance of common pipistrelle écholocation calls using environmental variables, **i) hypothesizing that the bat pass abundance depends on the vegetation spatial heterogeneity and on the area and height of vegetation and buildings, at different spatial scales.** In a second step, we estimate habitat availability for the common pipistrelle by using the model to predict the pass abundance for the “All green areas” and “No private green areas” scenarios. We evaluate the contribution of private green area to habitat availability for the common pipistrelle by comparing the habitat availability values obtained under the two scenarios. Because of their large size, **ii) we hypothesize that public green areas are more important contributors to the availability of habitat area for the common pipistrelle,** and only marginally complemented by private green areas. In a third step, we estimate the habitat connectivity associated with habitat availability under the two scenarios using a circuit modeling approach. We evaluate the contribution of private green area to habitat connectivity of the common pipistrelle by comparing the connectivity values obtained under the two scenarios. Because of their scattered spatial configuration, **iii) we hypothesize private green areas to be the main providers of habitat connectivity in the city,** providing the bats with stepping-stone connectivity across the urban matrix between larger habitat patches centered on public green areas.

**Figure 1:**
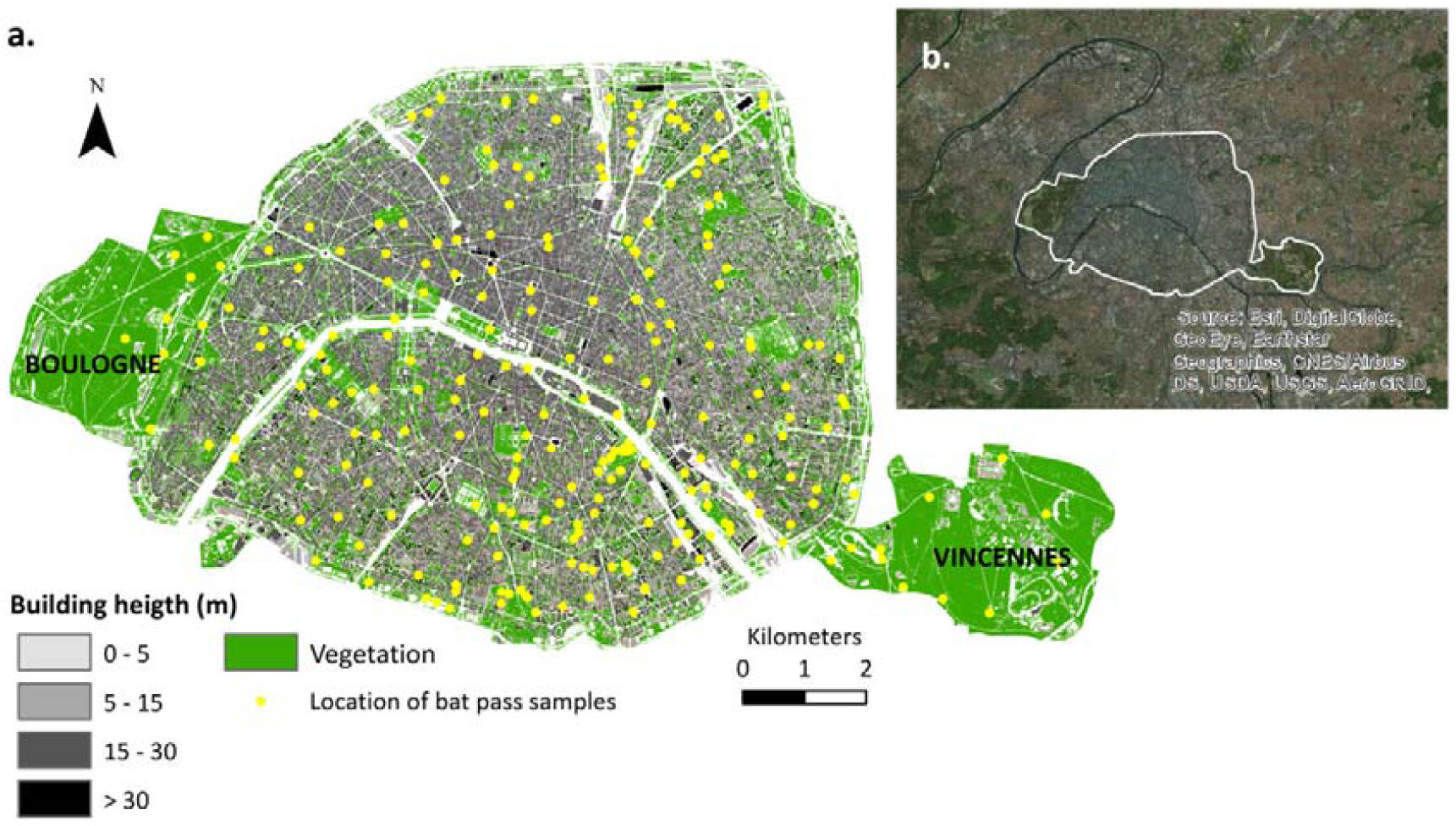
Map of the study area, i.e. intramural Paris. a. shows the location of the bat pass acoustic samplings used in the study and the two large parks (Boulognes and Vincennes) identified as the two habitat patches for the focal species *P.pipistrellus;* b. shows intramural Paris in its densified urban context, at the centre of the Paris megalopolis.

## 2. Methods

### 2.1. Study area

The study was conducted in “intramural Paris”, a densely populated city of 105 km^2^ (21,067 inhab/km^2^ in 2014) in the heart of the Greater Paris region. “Intramural Paris” refers to the central part of this agglomeration, bounded by the *Périphérique* ring-road, and administered by the *Mairie de Paris* (city council). Built areas are dominated by low-rise buildings of six to seven floors (i.e., 18 to 30 meters high) (Vincent Pellissier et al., 2012). The number and size of green areas is low compared to most other European big cities. Two woods – Vincennes (9.95 km^2^, east of Paris), and Boulogne (8.5 km^2^, west of Paris) – are the largest green areas in the city, bringing some nature right into the heart of Paris (Figure 2).

**Figure 2:**
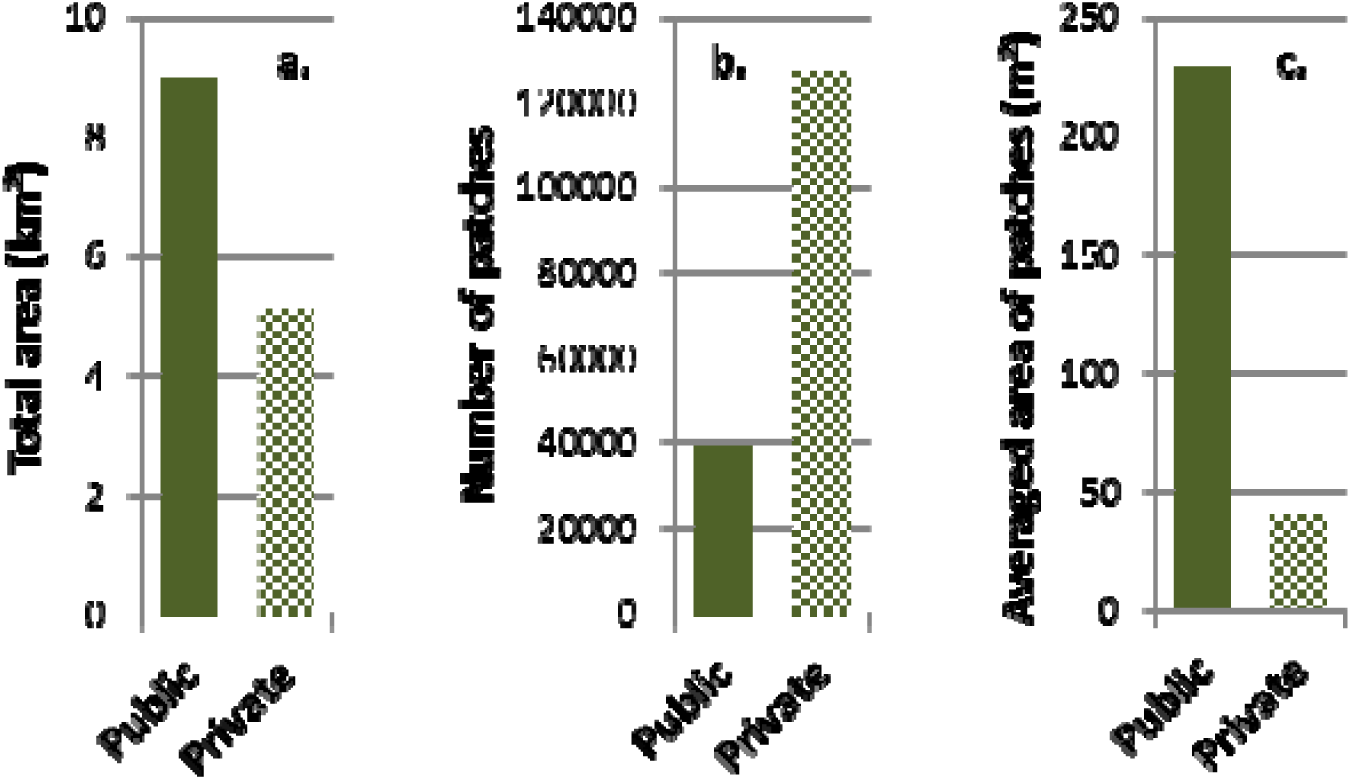
Comparison of public and private green areas in the Paris area using three basic landscape metrics: a. total area, b. total number of patches and c. average area of patches. The metrics have been computed for intramural Paris, excluding the Parks of Vincennes and Boulogne.

### 2.2. Step 1: Modeling and predicting bat pass abundance in Paris

#### 2.2.1 Bat sampling

We produced habitat availability and connectivity maps for *P. pipistrellus* using data from the French Bat Monitoring Programme (FBMP) for 2008 to 2013. The FBMP is coordinated by the French National Museum of Natural History and follows a standardized data recording methodology (Kerbiriou et al., 2018) (Appendix S1). The sampling scheme consists of randomly chosen 2×2 km squares from a 2×2 km grid. Within each square, observers selected and visited 10 points: at least five of these points were representative of the habitats of the square and the others were located in ‘favorable’ places for bats such as along the edge of woods. Each point was sampled using a continuous recording of 6 minutes and the ten points of a site were sampled on the same night and always in the same order at each visit. Observers recorded bats only when weather conditions were favorable (i.e., no rain, temperature higher than 12°C and wind speed of less than 5 m/sec; Appendix S1). Observers conducted the sampling during peak daily activity, i.e. beginning thirty minutes after dusk (FBMP recommendations). It generally takes less than 3 hours to sample the 10 recording points of the square (Vandevelde, Bouhours, Julien, Couvet, & Kerbiriou, 2014). Data was recorded from 2008 to 2013, between the 16^th^ June and the 5^th^ August.

We extracted the 224 intramural Parisian points from the FBMP database for 2008 to 2013 (Figure 1). Depending on observers’ availability over the years, the sampling effort at each point varied from one to five years. The dataset included 552 recordings at 224 points (139 points in 2008, 158 points in 2009, 72 points in 2010, 30 points in 2011, 68 points in 2012, 77 points in 2013).

We cannot distinguish individual bats from their echolocation calls, making it impossible to calculate absolute bat density, so we used the number of bat passes recorded every 6 minutes as a measure of bat activity (for more detail see Appendix S1). A bat pass is defined as one or more bat echolocation calls during a sound recording of 1.2 s at x 10 time expansion (see Appendix S1 for the methodology used to detect the number of bat passes). The duration of the recording (1.2 s) is predefined by the ultra-sound detector (Tranquility Transect; David Bale, Courtpan. Design Ltd, Cheltenham, UK) (Roche et al., 2011). Bat activity is associated with greater biomass of prey (Ciechanowski, Zając, Bitas, & Dunajski, 2007; Tibbels & Kurta, 2003; Verboom & Spoelstra, 1999). Bat activity is used as a proxy for bat abundance in research literature (Kerbiriou et al., 2018; Newson, Evans, & Gillings, 2015) and strongly correlated with minimum bat abundance in our data set (Appendix S1). Bat activity has shown its usefulness for studying anthropic impacts on bats (Millon, Julien, Julliard, & Kerbiriou, 2015; Wickramasinghe, Harris, Jones, & Vaughan, 2003), and for estimating habitat suitability in terms of food resources and accessibility (Frey-Ehrenbold, Bontadina, Arlettaz, & Obrist, 2013; Pinaud, Claireau, Leuchtmann, & Kerbiriou, 2018; Raino, 2007; Russo & Jones, 2003).

We recorded many surveys having 0 passes and few surveys having over 4 passes (4%). In order to limit over-dispersion in statistical analysis, we thresholded the maximum pass abundance at 4, meaning that recordings with more than 4 bat passes were attributed a number of 4. For all analyses, we averaged the abundance of bats passes observed per sample point over the different years.

#### 2.2.2. Creating variables for built areas and vegetation

We used a set of 18 variables to describe the built areas and the vegetation around each pixel. We chose variables known to influence the probability of observing the common pipistrelle because of their power to indicate resource availability, nesting opportunities, movement facilitation, or avoidance behaviour. Each variable was computed for three radii around the pixel, i.e., 20 m, 200 m and 500 m, to account for local to landscape-scale processes. 20 m corresponds to the minimum detection distance for the echolocation signal of search flight (i.e. echolocation calls before prey detection) (Barataud, 2015; Kalko & Schnitzler, 1993), while 500m corresponds to bat home range during the reproduction period (Davidson-Watts, Walls, & Jones, 2006).

##### Built areas and vegetation data

*APUR (Agence Parisienne d*’*URbanisme*: Parisian Urban Planning Agency) provided the data on building and vegetation location and height for the year 2012. The data were prepared by *APUR* based on several orthophoto images with a resolution of 0.5 m that we aggregated to a resolution of 2m. The data and their metadata containing more detailed information about data preparation are freely downloadable from the *APUR*’*s* website (Atelier Parisien d’Urbanisme, 2014; Atelier Parisien D’urbanisme, 2016). We also obtainws the location of the public green areas from the *APUR* website (Atelier Parisien d’Urbanisme, 2016). The location of the private green areas was estimated by subtracting the vegetation of public green areas from overall vegetation. In order to provide a broad overview of the respective spatial organization of public and private green areas in Paris, we computed three simple landscape metrics from the raw data: total area, total number of patches and average area of patches, a patch being defined as a unit composed of adjacent pixels of vegetation. For this simple descriptive analysis, a patch of green area was defined as continuous cells covered by vegetation.

##### Variables describing the vegetation

We computed four different variables based on vegetation height to describe the vegetation environment of the pixel. *P. pipistrellus* is known to be sensitive to vegetation: it typically commutes at a height of ∼3–10 m (Berthinussen & Altringham, 2012; Verboom & Spoelstra, 1999); it tends to avoid open habitats and vegetation higher than 3m mitigates the negative effects of urbanization (Hale et al., 2012). We therefore classified vegetation height into three classes: (i) <1m, (ii) 1 to 3m and (iii) >3m. We computed the total area covered by these three classes of vegetation and estimated a fourth variable, i.e., the spatial heterogeneity of the height of the vegetation around each pixel, using the standard deviation of vegetation height. We computed these four variables for the three radii (20 m, 200 m and 500 m radius), resulting in 12 vegetation variables in total. We calculated these 12 variables accounting for all green areas in the “All green areas” scenario, and repeated this process for the “No private green areas” scenario, only accounting for vegetation located in the public green areas (i.e. excluding the vegetation located outside of the public green areas).

##### Variables describing built areas

Buildings can be a barrier to movement (Hale et al. 2015) but may also be used for roosting (Simon et al., 2004). Furthermore, intermediate building height has been shown to be linked to higher abundance of insectivorous bird species in Paris (Vincent Pellissier et al., 2012). Beyond their direct effects on movement and habitat availability, buildings’ height and density can also be considered as a more general indicator of anthropogenic pressure, correlating with light and noise disturbance on an urbanization gradient (Grimm et al., 2008). We classified buildings within two height classes. Buildings under 15 m mainly consisted of low-rise buildings and individual houses. This class of building was dominant in the external districts of Paris (Figure 1). Buildings of over 15m were mainly located in the old center of Paris and in the north-west. For each of the three scales detailed earlier (i.e., 20 m, 200 m and 500 m radius), we computed the area covered by the two building classes and attributed the value to the central pixel.

#### 2.2.3. Modeling bat pass abundance using vegetation and built areas variables

We modeled the relative pass abundance of P. *pipistrellus* with the previously described vegetation (12 variables) and built areas (six variables), using a boosted regression tree modeling approach (gbm) (gbm package; Greg Ridgeway with contributions from others, 2017) using R 3.4.0 (R Core Development Team, 2018). The gbm approach was relevant in this study because it can handle a large number of predictors – even collinear ones – and deal with spatial autocorrelation effectively, and it also has a strong predictive performance (Elith, Leathwick, & Hastie, 2008). Because the data contained a lot of zeros (130 points out of a total of 224 points), we employed a Hurdle modeling approach to account for zero-inflation (Potts & Elith, 2006; Povak et al., 2013). This approach is consistent with previous studies modeling bat passes (Aurelie Lacoeuilhe, Machon, Julien, Le Bocq, & Kerbiriou, 2014; Vandevelde et al., 2014). The Hurdle model is a two-step modeling approach. The first model, run on data transformed into Presence /Absence pass data, calculates the probability of pass occurrence. The second model is fitted on the pass abundance data (excluding absence data), and aims to predict pass abundance only where bat passes are predicted to potentially occur in the first model. When used for prediction purpose on a new set of environmental data, the Presence/ Absence model is run first. If the predicted value is below a fixed pass occurrence probability (assimilated to predicted absence), the predicted value is maintained. If the predicted value is above the fixed threshold, then it is the value predicted by the abundance model that is retained. Based on the results of Presence/Absence modeling, we fixed the threshold between absence and presence at 0.45. Because the number of years of observation varied between the different points, we weighted the points by the number of years of observation. We calibrated the models with a learning rate of 0.0005, an interaction depth of 4 (meaning that we consider interactions), a minimal number of individuals per leaf of 10, and a fraction of 0.6 for training the algorithm (Elith et al., 2008). We checked for and did not find any significant residual spatial autocorrelation.

### 2.3. Step 2: Evaluating the contribution of private green areas to foraging-commuting habitat availability

#### 2.3.1. Estimating foraging-commuting habitat availability

We used the fitted Hurdle model to predict the bat pass abundances over the entire study area for the “No private green areas” and the “All green areas” scenarios. We then used the resulting bat pass abundance as foraging-commuting habitat availability maps. For the “All green areas” scenario, predictions were based on the vegetation variables encompassing all green areas, whereas for the “No private green areas” scenario, vegetation variables were restricted to the green public areas. We measured total foraging-commuting habitat availability (hereafter referred to simply as habitat availability) for each scenario as the sum of all predicted pass abundance in intramural Paris.

#### 2.3.2. Contribution of private green areas to foraging-commuting habitat availability

We estimated the contribution of private green areas to habitat availability by subtracting the total predicted pass abundance of P. *pipistrellus* over the entire study area of the “No private green areas” scenario from the predicted pass abundance for the “All green areas” scenario.

### 2.4. Step 3: Evaluating the contribution of private green areas to foraging-commuting habitat connectivity

#### 2.4.1. Conductance maps

Conductance maps depict the ease of movement across the mapped area that varies for example with land cover, habitat quality or slope. Conductance maps are used as input data to model connectivity under the Circuit approach. We used the habitat availability maps as conductance maps. This data-based approach is expected to produce more realistic conductance values than would be obtained using expert opinion.

We changed the resolution of the two habitat availability maps from 2m to 20m by averaging the values of the cells. This change in resolution was needed to be coherent with the bat’s perceptual grain, defined as the grain at which an organism responds to the heterogeneity of the landscape (Wade, Mckelvey, & Schwartz, 2015; Wiens & Milne, 1989) (see Appendix S2 for details). As P. *pipistrellus* can only detect its prey within a maximum radius of 3.5 m for small prey and of 15m for large prey (Holderied & Helversen, 2003), we consider a resolution of 20m would adequate fit its perceptual range.We then summed predicted bat passes within all pixels for intramural Paris (excluding the Vincennes and Boulogne parks) to obtain a simple indicator of habitat availability. We obtained the conductance maps by rescaling the predicted pass abundances to values between 0 and 10,000.

#### 2.4.2. Building connectivity maps

We identified two areas, i.e., Boulogne and Vincennes parks, as the two source habitat patches of the common pipistrelle and we looked for connectivity paths between them (Figure 1). These parks are the two largest green areas of Paris, and are known to house viable populations of P. *pipistrellus.* Based on our data, bat activity in these parks (2.7±0.9 bat passes/6minutes on average) is greater than in intramural Paris by a factor of 2.1. These parks are located on opposite sides of Paris and this makes them ideal for investigating how and where bats pass through the city matrix.

We used Circuitscape, a program that uses circuit theory to model connectivity in heterogeneous landscapes, to assess landscape connectivity (McRae & Beier, 2007). The Circuit theory approach provides a continuous estimate of connectivity within the area studied, i.e., for each pixel, integrating all possible pathways. The results provided by Circuitscape include connectivity maps based on conductance and a measure of the total conductance between the two habitat patches (i.e. Vincennes and Boulogne parks).

#### 2.4.2. Contribution of private green areas to foraging-commuting habitat connectivity

We estimated the contribution of private green areas to habitat connectivity by subtracting the total conductance value provided by Circuitscape for the “No private green areas” scenario from the total conductance value for the “All green areas” scenario - for the entire study area.

## 3. Results

### 3.1. Spatial characteristics of public and private green areas in Paris

The total green area belonging to private owners was smaller than the total public green area (36.4% and 63.6 %, respectively; Figure 2a). The private green areas were much more fragmented, as illustrated by a number of patches that was three times higher than for public areas and an average patch size five times smaller (Figure 2b and 2c).

### 3.2. Step 1: Modeling and predicting bat pass abundance in Paris

The gbm analyses produced contrasting results for Presence /Absence and abundance of passes data for the built areas and vegetation variables and their scale of impact, suggesting different processes underpinning pass occurrence and abundance (Figure 3). The Presence/Absence of bat passes was dependent on large-scale environmental conditions (200m to 500m radius). The passes were more likely to occur in areas with a high proportion of vegetation and less likely to occur in areas with large concentrations of buildings under 15m high. The occurrence of bat passes revealed a higher abundance of bat passes when buildings are smaller (under 15m high), at 200m to 500m radius. The abundance of passes was mainly driven by local conditions (20 m) and to a lesser extent by larger-scale conditions (200 and 500 m). Locally, the high proportion of tall buildings was the strongest driver, negatively impacting the bat pass abundance. Overall, the proportion of buildings tended to decrease the abundance of passes at different scales. Conversely, a large proportion of tall vegetation as well as the variation in vegetation height at local and medium scales (20 and 200m) tended to be beneficial (Figure 4).

**Figure 4:**
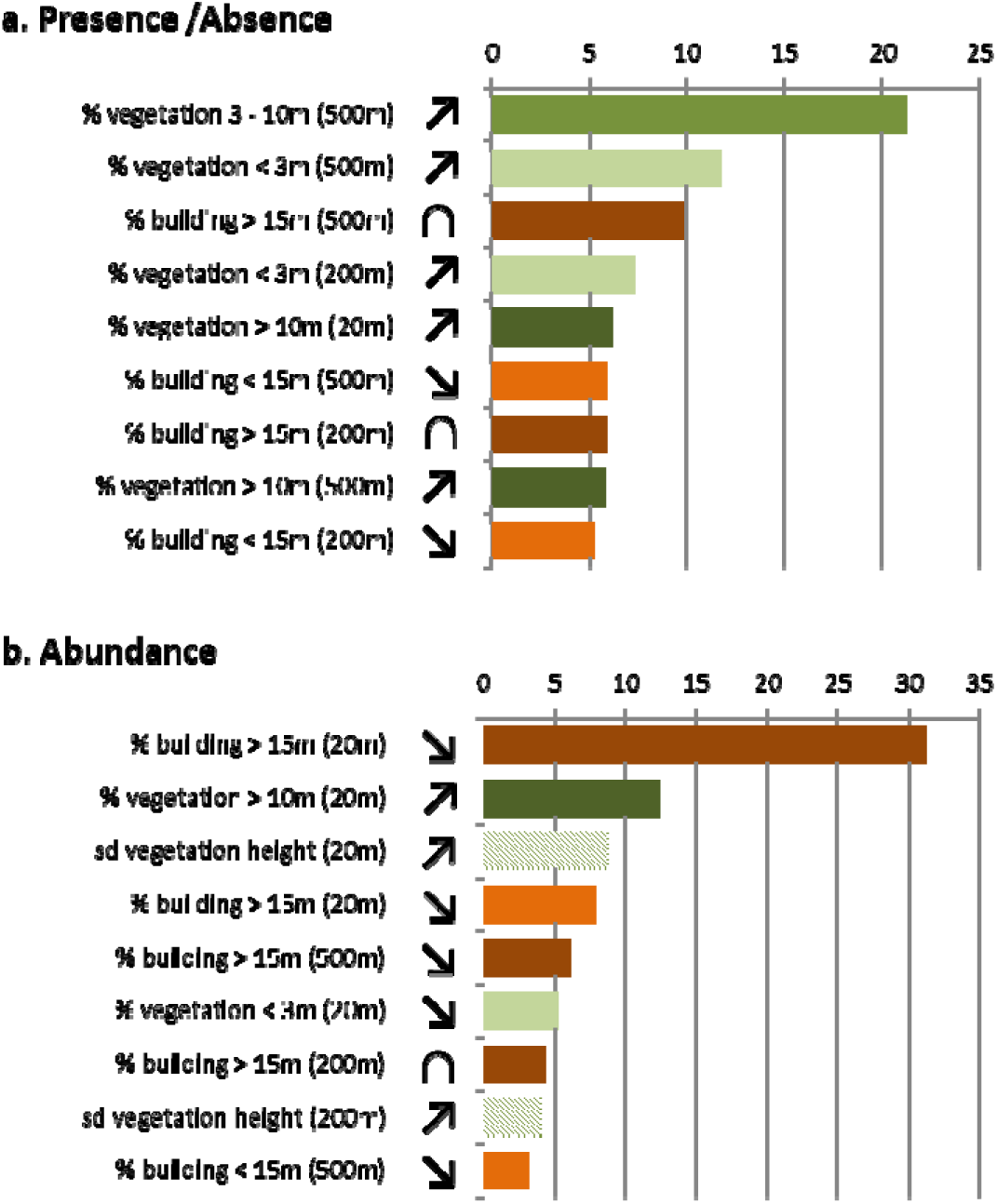
Variables ordered by their contribution (in percent) to the occurrence and bat pass abundance as estimated by the two gbm analyses, a. the Presence /Absence data, and b. the abundance of passes data. The contribution is the relative influence of each variable on the response variable. The signs indicate the general form of the response of the bat pass abundance to each variable, i.e., increasing, decreasing or unimodal. Only the variables with higher contributions are shown.

### 3.3. Step 3: Evaluating the contribution of private green areas to foraging-commuting habitat availability

The interpolated predictions of bat pass abundance for the “All green areas” scenario, showed that larger predicted pass abundances were mainly concentrated in the two large parks of Boulogne and Vincennes (Figure 5) and to a lesser extent within Paris’ larger parks. In intramural Paris, the highest predicted pass abundances were found mostly in the southern and eastern peripheral areas. The map of predicted pass abundances under the “No private green areas” scenario also identified the two large parks as main areas of habitat (Figure 5). The areas with higher predicted habitat availability were similar in the two scenarios, the main difference between layd in the predicted bat pass abundance, which was much lower in the “No private green areas” scenario (Figure 5). The predicted pass abundances were extremely low in the dense city center, north of the Seine River. We used the total number of predicted bat passes in intramural Paris (excluding the two woods) as indicators of the area of available habitat. The total number of predicted bat passes was 12,874,960 in the “All green areas” scenario, and 6,709,623 in the “No private green areas” scenario. The difference, i.w., 6,165,337 bat passes, corresponded to the contribution of private green areas to available habitat, and therefore represents 47.9% of the contribution of total green areas to available habitat while private green areas represent just 36.4% of the total green area.

**Figure 5:**
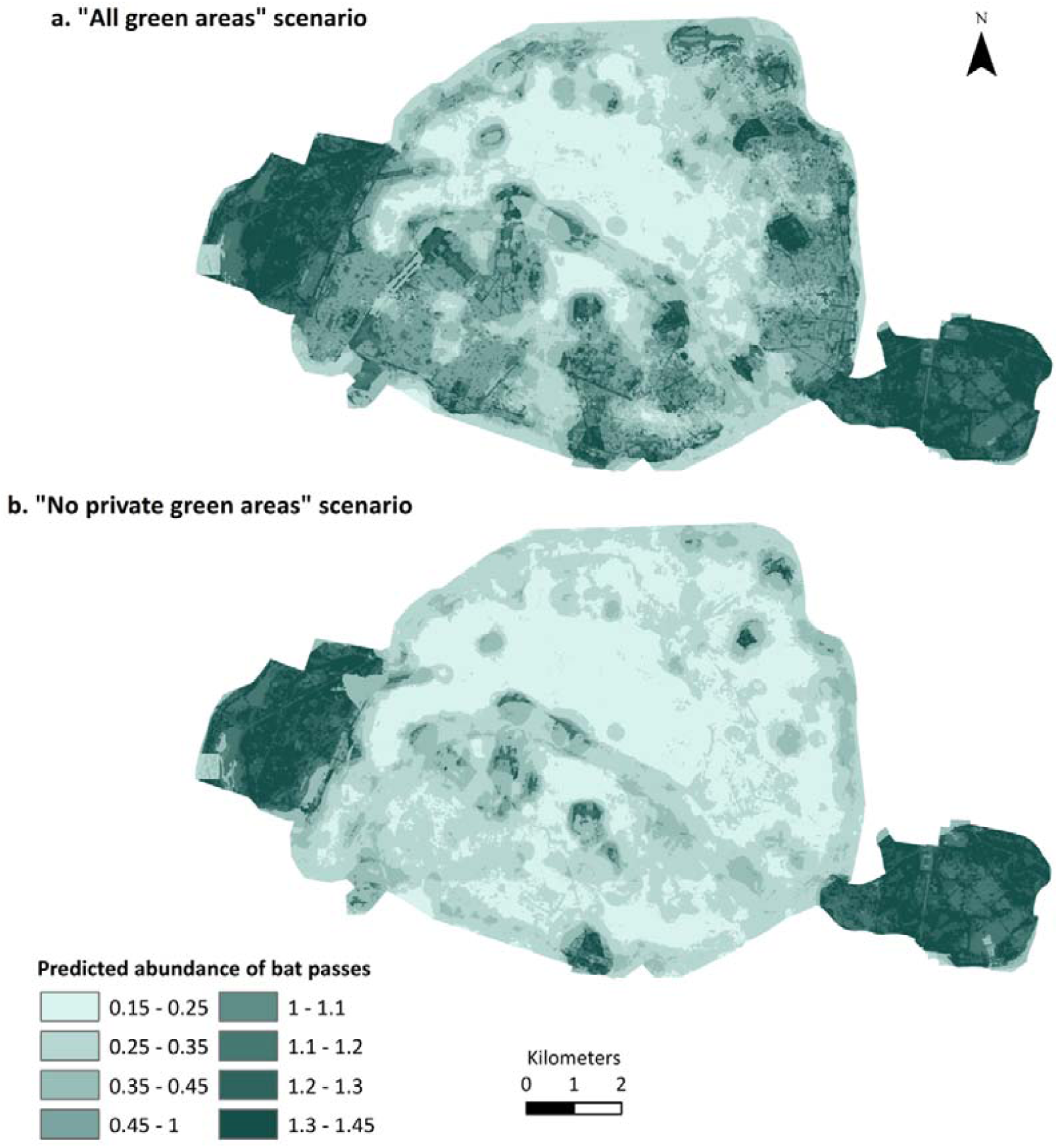
Habitat availability maps of *P. pipistrellus* in Paris showing the bat pass abundance as predicted by the gbm for a. the “all green areas” scenario and b. the “No private green areas” scenario.

### 3.4. Step 3: Evaluating the contribution of private green areas to foraging-commuting habitat connectivity

The spatial structuring of the conductance maps under the two scenarios revealed close spatial structures with connectivity paths mainly located in the south of Paris (Figure 6). However, the total resistance of intramural Paris was estimated to be 0.059 in the “All green areas” scenario, compared to 0.112 for the “No private green areas” scenario, meaning that private green areas decreased the city’s total resistance by 88.7% when compared to the resistance of the “No private green areas” scenario.

**Figure 6:**
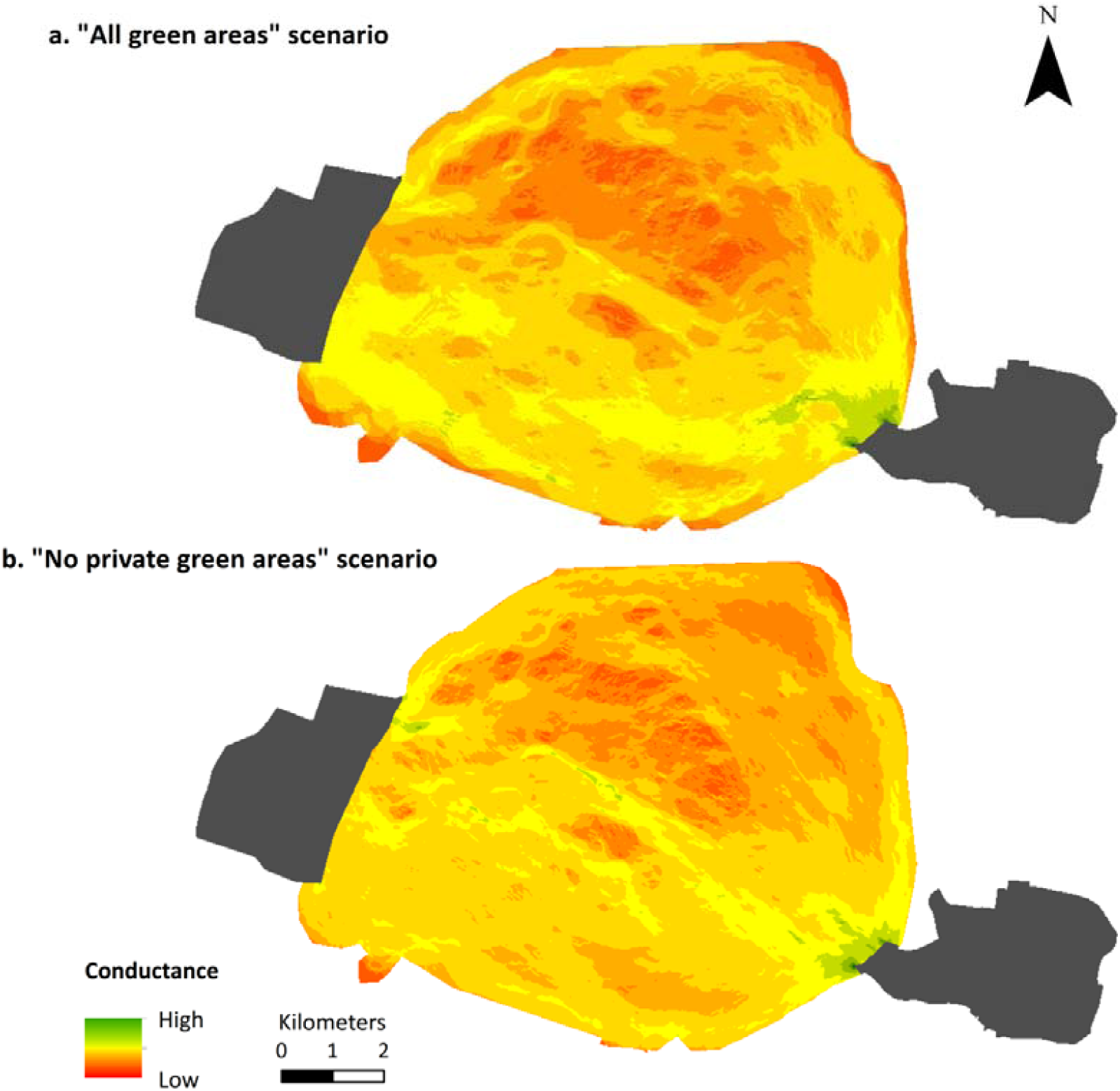
Conductance maps for *P. pipistrellus* in Paris linking Boulogne park (left black patch) and Vincennes park (right black patch) for a. the “all green areas” scenario and b. the “No private green areas” scenario.

The connectivity map for the “All green areas” scenario shows stronger connectivity paths passing through the peripheral areas of the city with the southern area showing higher connectivity levels. In the “No private green areas” scenario, the Seine River appeared as the preferred path across Paris, concentrating a large proportion of the flow. The contribution of private green areas to connectivity did not follow clear spatial patterns, either creating new paths or strengthening exiting paths between public green areas, especially in the south (Figure 6).

## 4. Discussion

The aim of the present study was to estimate the contribution of private green areas to habitat availability and connectivity for the common pipistrelle in Paris. By focusing on this objective, our study tackles important questions regarding the functional role of urban green areas in a dense urban setting, i.e. the importance for the common pipistrelle of the total area of green space, vegetation heterogeneity, the structure of the urban matrix and the connectedness of green areas (Lepczyk et al., 2017; Shwartz et al., 2014). By proposing a number of pointers, the output of this study provides scientific elements supporting the idea that private green areas could be worthwhile targets for conservation strategies in cities (Lepczyk et al., 2017).

### 4.1. The importance of vegetation and built areas characteristics for foraging-commuting habitat availability for the common pipistrelle

*P. Pipistrellus* is among the more habitat generalist of bat species (Regnery, Couvet, Kubarek, Julien, & Kerbiriou, 2013) and is regularly found in urban areas (Bartonicka & Zukal, 2003; Hale et al., 2012; Vandevelde et al., 2014). In line with previous findings, we observed a positive impact on bat occurrence of the total area covered by vegetation at larger scales (200 and 500m) (Azam et al., 2015; Aurelie Lacoeuilhe, Machon, Julien, & Kerbiriou, 2016). However, while occurrence was more effectively predicted by larger scale environmental conditions (200m to 500m), bat pass abundance was largely driven by local conditions (20m). In other words, while the probability of bat presence was linked to conditions at broad scales, local-scale conditions were predominant in explaining the location of the paths At such local scales and in line with previous findings, our results showed that higher bat pass abundance is linked to a selection of woody habitats and the avoidance of open habitats (Bartonicka & Zukal, 2003; Hale et al., 2012; C. G. Threlfall, Williams, Hahs, & Livesley, 2016) and to variation in vegetation height (Suarez-Rubio, Ille, & Bruckner, 2018). The proportion of built areas had a negative overall impact on the presence and abundance of passes although an intermediate proportion of higher buildings appeared to be beneficial for the species (Hale et al., 2012). A similar high building effect has previously been observed in Paris for insectivorous birds, so we may hypothesize that this building structure benefits insectivorous species’ foraging activity possibly by concentrating the insects in certain areas (Vincent Pellissier et al., 2012).

### 4.2. The contribution of private green areas to the foraging-commuting habitat availability of the common pipistrelle

Higher bat activity was recorded and predicted in the peripheral areas of Paris where the higher density of private green areas both enhanced the attractiveness of large public green areas and greatly extended their benefits for the species to the surrounding areas of the city via a net buffering effect. These results highlight the complementary contribution of private and public green areas to habitat availability and quality, a process known as “Ecological land-use complementation” in research literature (Colding, 2007). Here, we observed that large public green areas constitute the main patches of available habitat in the city (core areas) while private green areas increase their capacity to support individuals and enlarge their effective area. Comparable complementation effects of gardens for public areas have been documented elsewhere While the importance of private green areas for the common pipistrelle has already been demonstrated in previous studies (Hale et al., 2012), here we have also shown that in Paris private green areas have a disproportionately positive impact on habitat availability *vis-à-vis* their total coverage. Thus, while private green areas only represented 36.4% of the total green area, we found that they actually supported 47.9% of total habitat availability (i.e. foraging-commuting activity) for the common pipistrelle. This importance could be attributable to the differential types of vegetation favored in private green areas when compared with public green areas. Thus, the areas with high density of private green areas also appear to have higher availability of taller vegetation, which is an important driver of bat activity in our study.

### 4.3. The contribution of private green areas to foraging-commuting habitat connectivity for the common pipistrelle

The complementation effect between private and public green areas for the common pipistrelle was even stronger in the case of connectivity, as private green areas decreased the total resistance of the city by 88.7% even though they only represented 36.5% of the total green area. Thus, the spatial configuration of private green areas in the city appeared to be very important for the common pipistrelle, providing the stepping stones between the public green areas that serve as the nodes of the urban network (Rudd et al., 2002). If private green areas consist of small patches uniformly distributed across the city, they appear fragmented but not isolated, with the notable exception of the city center which appears highly resistant for the species. A study focusing on the role of green areas at business sites in the Parisian ecological network drew comparable conclusions, enhancing the functional connectivity role of green areas at business sites as stepping stones (Serret et al., 2014).

### 4.4. Private green areas for conservation

Our findings demonstrating the complementary role of public and private green areas for habitat availability and connectivity tend to confirm the benefits of moving towards more inclusive conservation strategies that include both public and private stakeholders in cities (Rands et al., 2010). Private areas have been found to contain more plant diversity and rarer species than public areas (Politi Bertoncini et al., 2012). “Wildlife-friendly gardening”, as time investment in the garden and reducing pesticide, use have been shown to positively impact wildlife (Goddard et al., 2013; Muratet & Fontaine, 2015). The challenge for conservation is therefore to organize and manage these “human-occupied” areas to increase their ecological value (Dearborn & Kark, 2010). Thus, it is the way in which private green areas are managed that could be targeted by new conservation initiatives in cities, following the ones applied to public green areas over the past few years (City of Dublin, 2016; City of London, 2016; Conseil de Paris, 2018; Ville de Paris, 2017).

### 4.6. Limits of the approach

Our approach has certain limitations. First, as we did not have information about the location of private green areas, we bypassed this problem by only considering public green areas and inferring the importance of private green areas by subtracting the values obtained for the “No private green areas” scenario from the “All green areas” scenario. This method can induce small biases if the delineation of public areas is not perfect, by artificially increasing the importance of private green areas by a small amount. However, because we focussed on the relative differences between the two scenarios in terms of total area, contribution to habitat availability and contribution to connectivity, the impact of such a distortion on the study outputs should be negligible. Second, we limited our study area to “intramural Paris”, excluding the surrounding urban areas which are a little less densified. In other words, we excluded potential connectivity paths linking the two parks but bypassing Paris. The existence of such paths would reduce the flux of individuals flying in/through Paris but would not change the observations concerning habitat availability and connectivity patterns.

## Appendices

### Appendix S1: Detailed information on recording settings and conditions

#### 1. Protocol

##### 1.1. General description

**Table S1-1:**
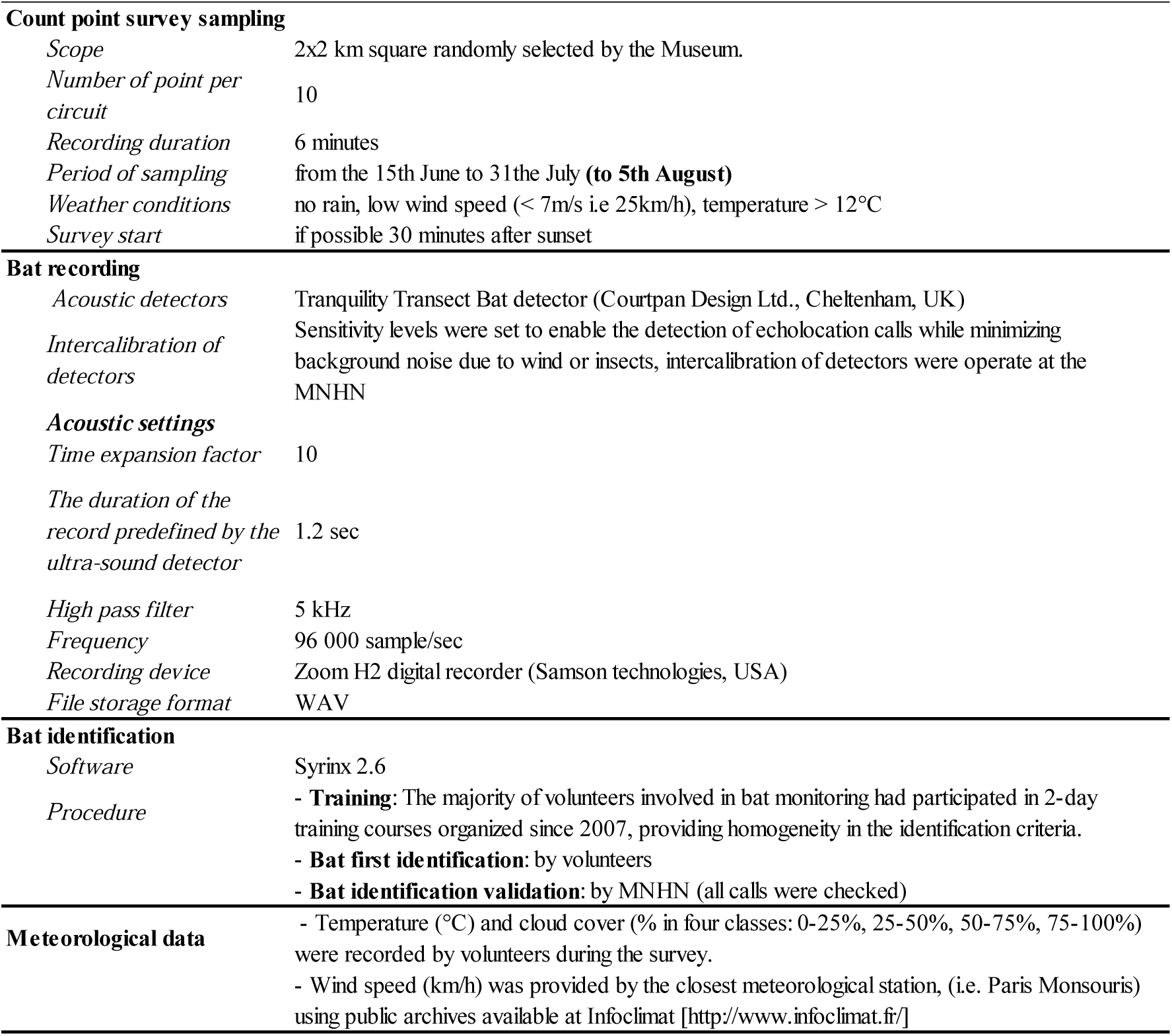
Characteristics of the protocol and sampling design used for the monitoring of the temporal trends of bat populations on a national scale by the French Bat Monitoring Programme (Vigie-Chiro, 2018), supervised by the French Museum of Natural History. In brackets and bold are indicated the small adaptations of the protocol made for the data used in the present study.

##### 1.2. Bat activity records

Records were done using Syrinx software version 2.6 (Burt, 2006) for spectrogram and Adobe Audition for spectral analysis together with Scan’R (Binary Acoustic Technology, 2010) to isolate each bat vocalization and automate measurement of relevant parameters (Gannon et al., 2004, Obrist et al., 2004, Barataud 2012). Sound species identification was verified by coordinators of the French Bat Monitoring Program. *P. pipistrellus* is a common bat which identification does not raise noticeable identification uncertainty, and even less in dense urban context such as Paris where species overlapping on acoustic repertoires do no occur.

##### 1.3. Measurement of bat activity

Bat activity at the point scale is calculated as the sum of the number of bat passes recorded per 6 minutes (Fig S1-2a). A pass of bat was defined as a call sequence containing one or more pulses and when the time between calls exceeded four times the inter-pulse interval (Parsons & Jones, 2000, Kerbiriou et al. 2019). In the protocol followed in this study, the duration of the record is fixed at 1.2 s. Within each 1.2 s record, the minimum number of bat recorded simultaneously was estimated based on inter-pulse interval and frequency (Fig S1-2b).

**Figure S1-2a:**
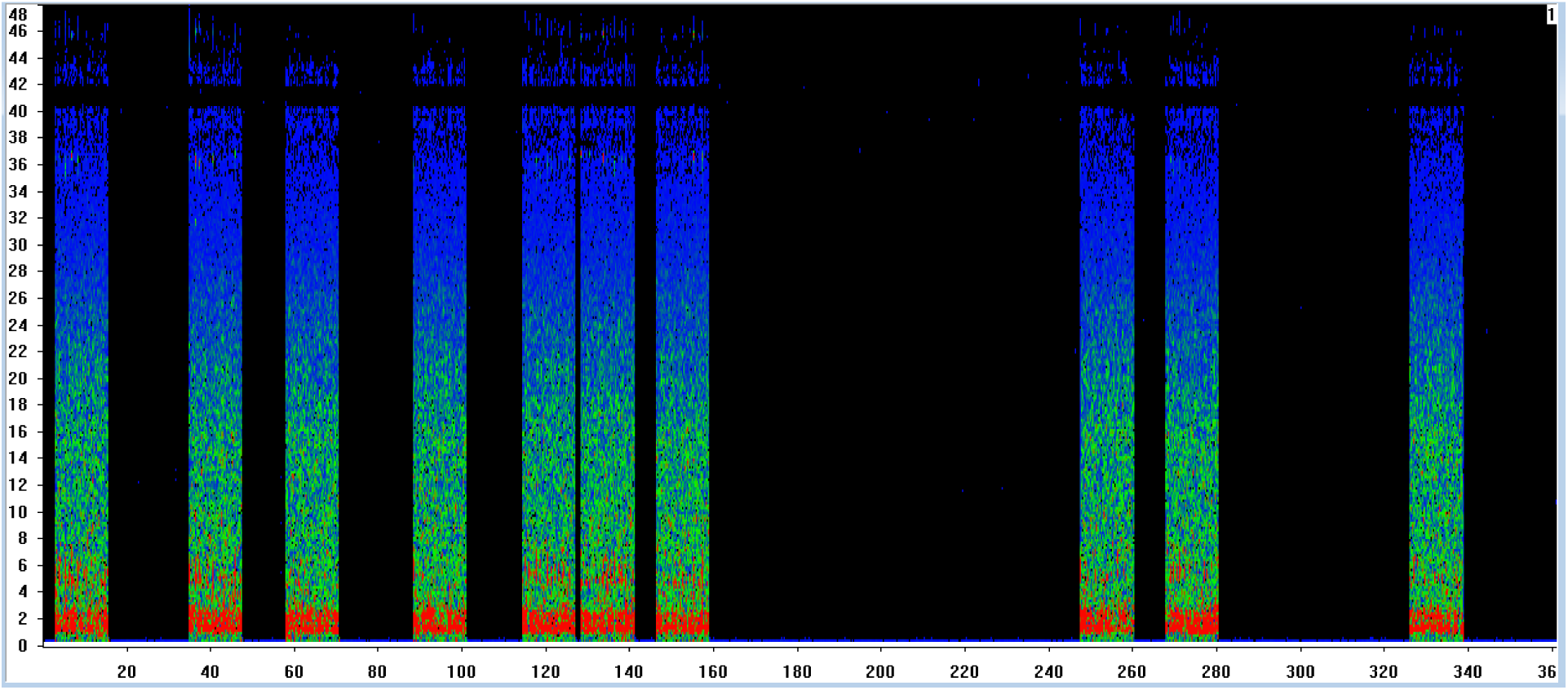
example of a 6 minutes record including 10 records of sound at a ×10 time expansion.

**Figure S1-2b:**
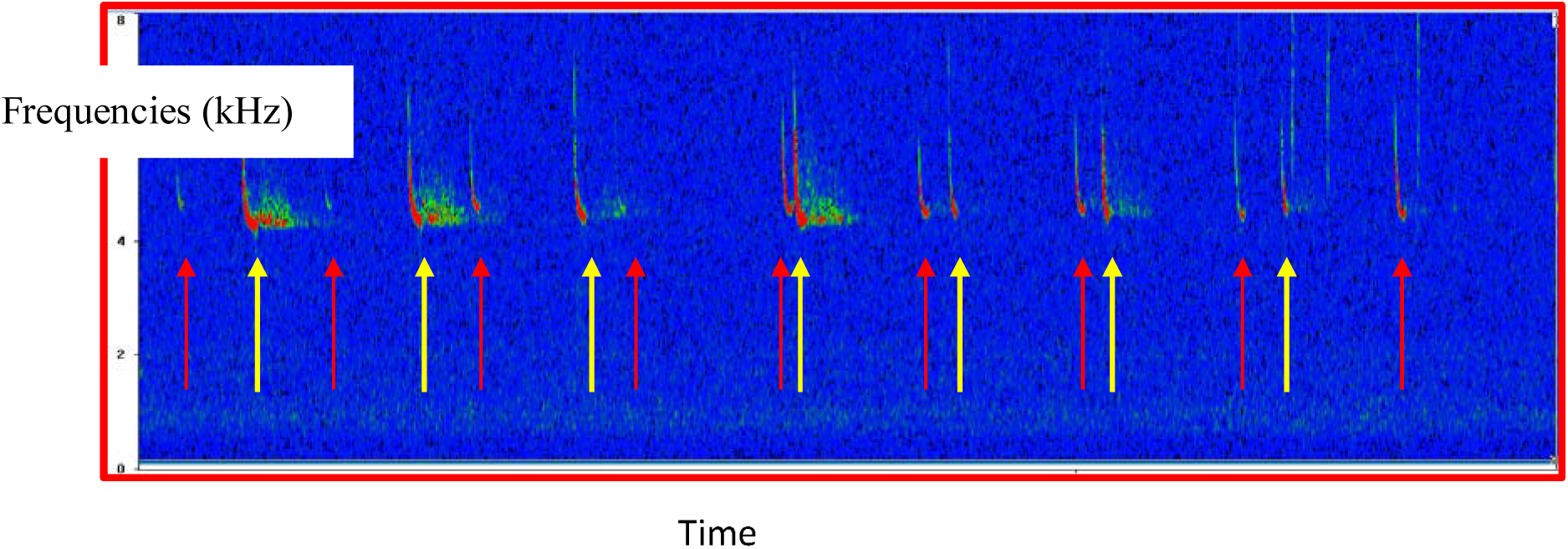
focus on a 1.2s sound sample (time expansion) recorded at time 130, which include 2 individuals of *P. pipistrellus* (highlight by red and yellow arrows).

##### 1.4. Relationship between bat activity and the number of contacted individuals

**Figure S1-3:**
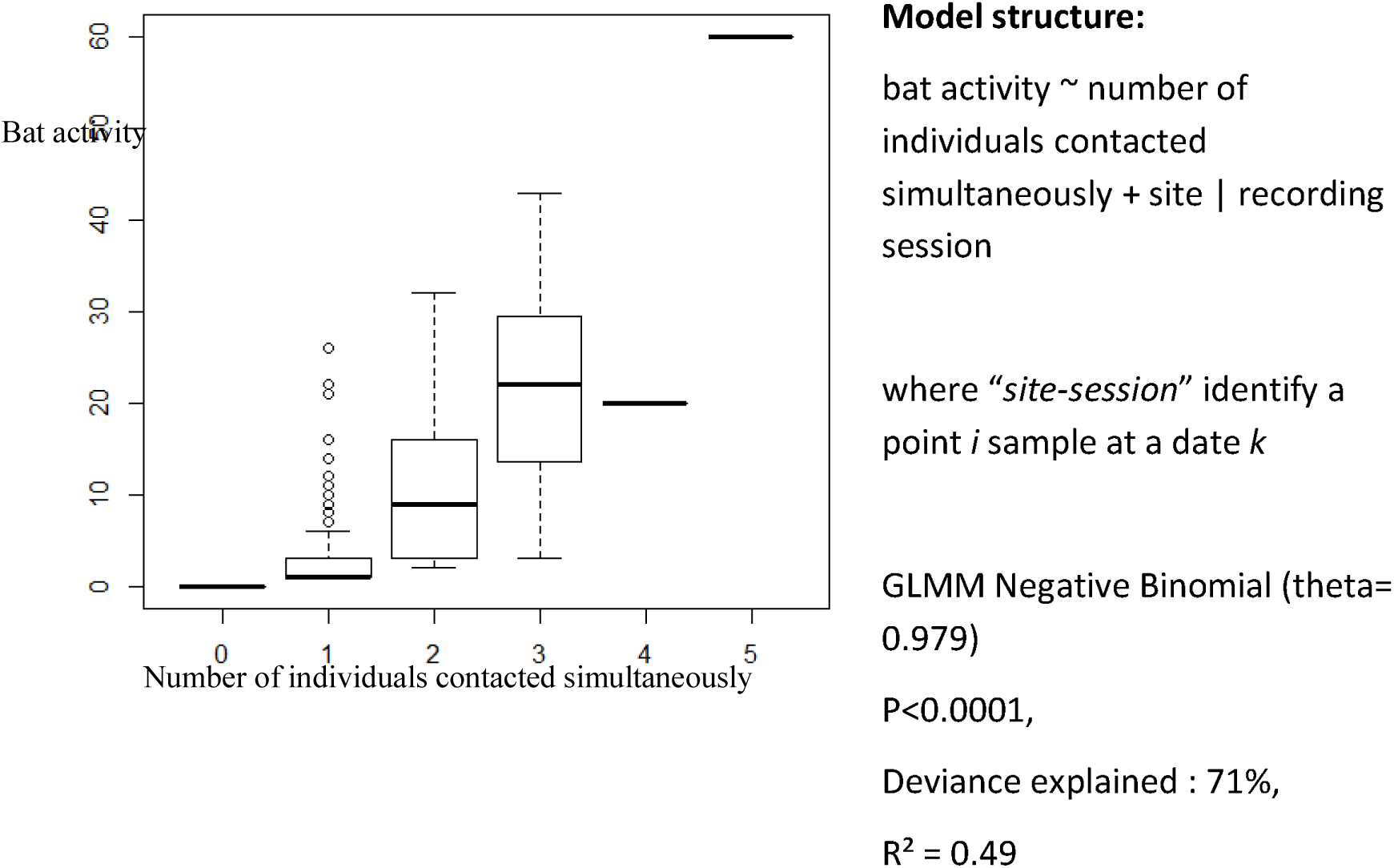
Relationship between bat activity and the number of individuals contacted simultaneously

###### Model structure

bat activity ∼ number of individuals contacted simultaneously + site | recording session

where *“site-session”* identify a point / sample at a date *k*

GLMM Negative Binomial (theta= 0.979) P<0.0001, Deviance explained: 71%, R^2^ = 0.49

### 2. Data sampling conditions

**Figure S1-1:**
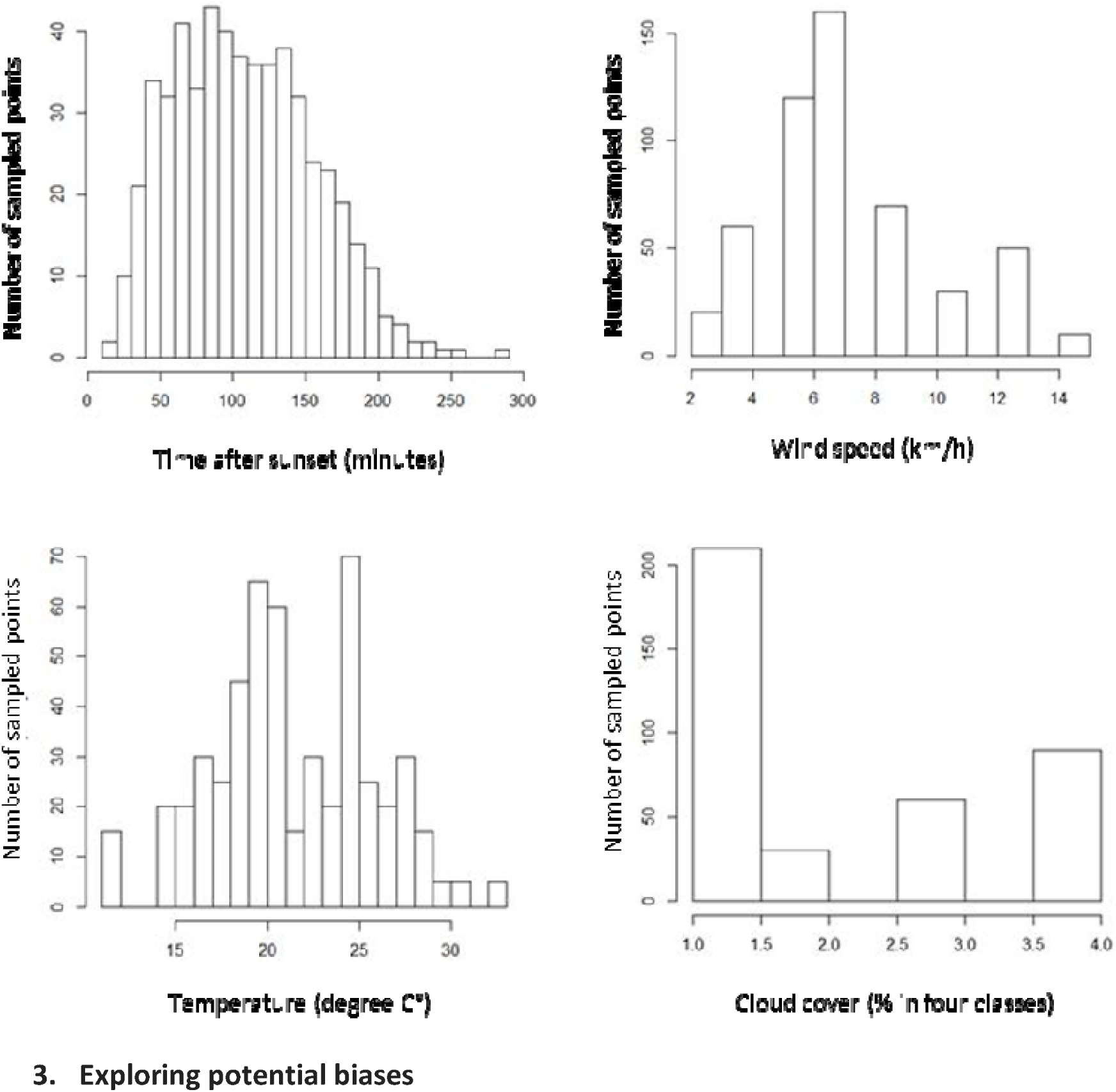
Time and weather sampling conditions

### 3. Exploring potential biases

#### 3.1. Patterns of nightly activity of P. pipistrellus

Volunteers are strongly encouraged to conduct their sampling during daily peak activity (Roche et al., 2005), i.e. beginning thirty minutes after dusk (FBMP recommendations). It usually takes less than 3 hours to sample the 10 recording points of each square: 97% of data collected in Paris occurred between 30 minutes after sunset and 3h30. During this period the pattern of nightly activity in urban context for *P. pipistrellus* is relatively flat (Fig. S1-4)

**Figure S1-4:**
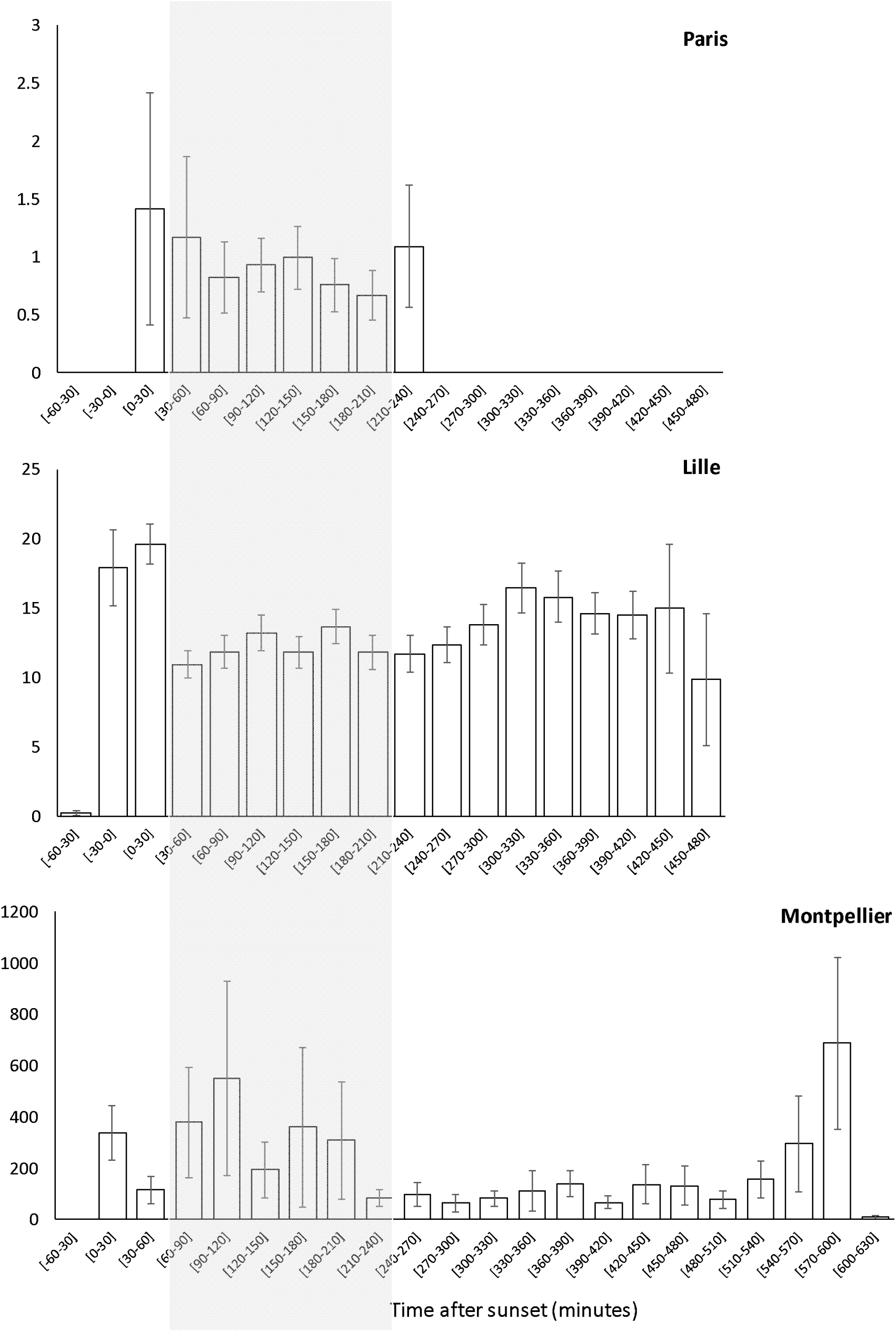
Pattern of nightly activity of P. *pipistrellus* in urban context Paris in three French cities, computed from the data of the French Bat Monitoring Programme: in Paris using the data used in the present study (552 6-minutes recordings, 224 points, years 2008-2013), Lille (n=73 full-nights recordings, 73 points, 2015; Pauwels *et al.,* 2019), and Montpellier (2012, 20 full-nights recordings, 20 points, unpublished data). The light grey section indicates the recording period recommend by the FBMP.

#### 3.2. Correlation between time of recording and the set of explanatory variables

**Table S1-2:** Correlation (Spearman’s *rho)* between time of recording and the explanatory variables

| Landscape-scale | 20 | 200 | 500 |
| --- | --- | --- | --- |
| *Buildings of over 15 m* | $\rho=0.048$ ; $P=0.265$ | $\rho=0.054$ ; $P=0.209$ | $\rho=0.027$ ; $P=0.522$ |
| *Buildings of under 15 m* | $\rho=0.055$ ; $P=0.200$ | $\rho=-0.070$ ; $P=0.1039$ | $\rho=-0.109$ ; $P=0.011$ |
| *Tree vegetation* | $\rho=0.029$ ; $P=0.503$ | $\rho=-0.017$ ; $P=0.690$ | $\rho=-0.023$ ; $P=0.593$ |
| *Shrub* | $\rho=-0.013$ ; $P=0.758$ | $\rho=-0.004$ ; $P=0.931$ | $\rho=-0.040$ ; $P=0.352$ |
| *Herbaceous* | $\rho=-0.014$ ; $P=0.743$ | $\rho=0.010$ ; $P=0.823$ | $\rho=-0.039$ ; $P=0.362$ |
| *Sd vegetation height* | $\rho=-0.042$ ; $P=0.325$ | $\rho=0.100$ ; $P=0.020$ | $\rho=0.082$ ; $P=0.056$ |

To protect from Type I error, a Bonferroni correction should be considered: for an alpha-value (α_*original*_=0.05) the Bonferroni correction (α_*adjusted*_ = α_*original*_/*k*; α_*adjusted*_=0.0028), thus to determine if any of the 18 correlations is statistically significant, the *P-value* must be *P*<0.0028

#### 3.3. Variation of explanatory variables among recording’s nights

**Figure S1-5:**
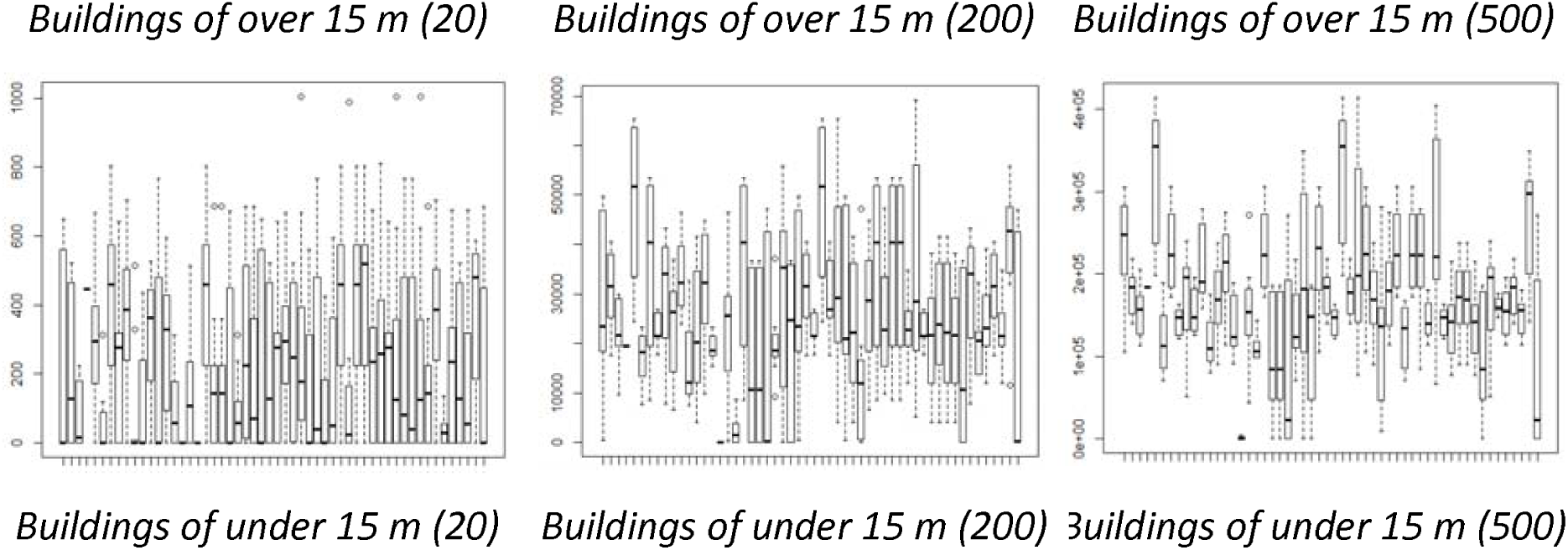
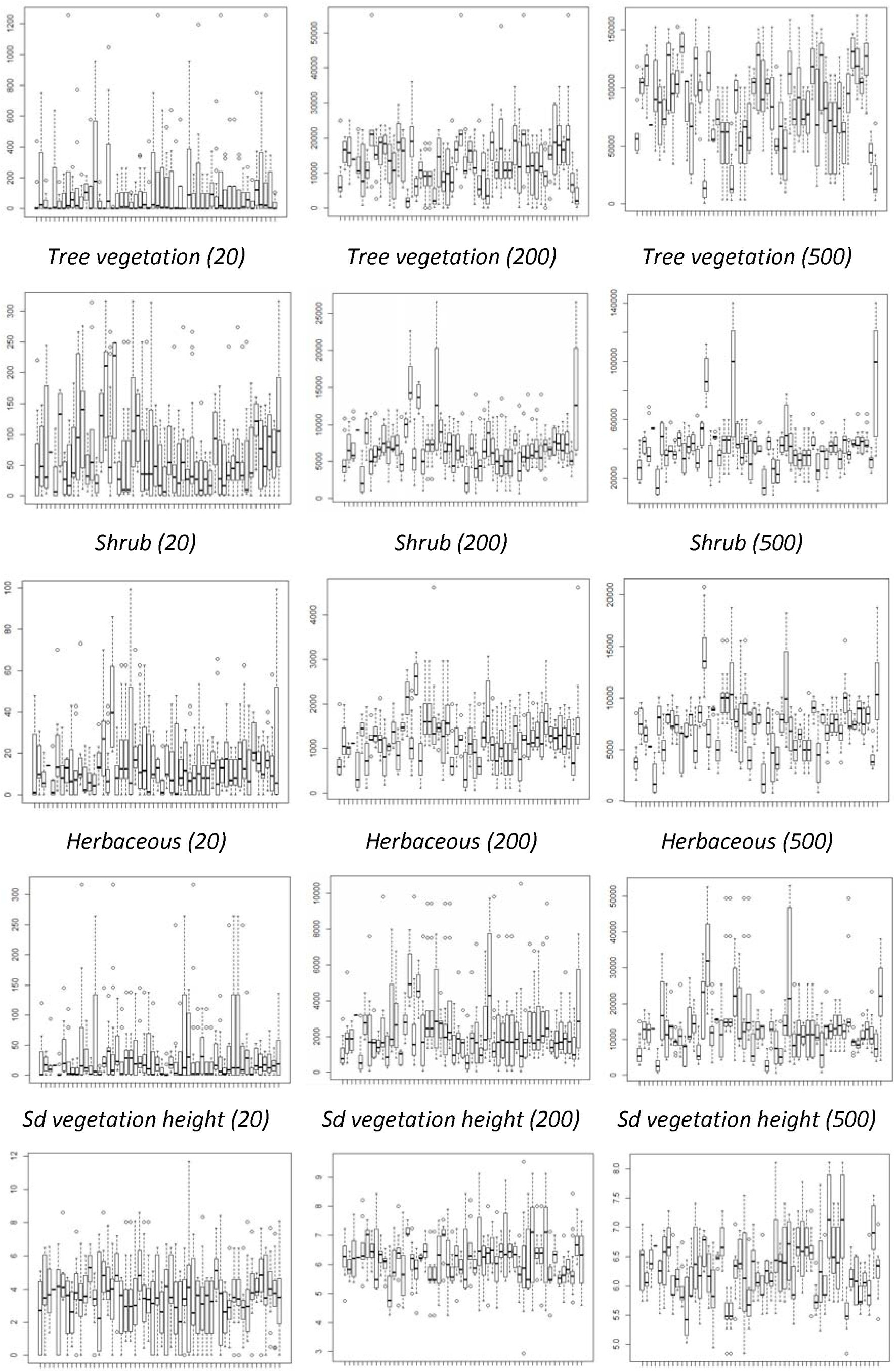
variations of explanatory variables among recording’s nights

## Appendix S2

Too small a resolution would result in increasing the resistance of the matrix by allowing very small elements (in this case 2m) to counter the movement, while the individual could in reality bypass the element.

We chose to aggregate data at this step, and not earlier in the analyses, mainly because the private green areas are usually small areas and would have disappeared with the degradation of the vegetation raster, resulting in an extensive loss of primary information for the object being studied. When tackled at this stage, the problem is much less important because we are dealing with continuous data that are simply averaged at the coarser resolution.

